# MitoTracer facilitates the identification of informative mitochondrial mutations for precise lineage reconstruction

**DOI:** 10.1101/2023.11.22.568285

**Authors:** Xuexin Yu, Jing Hu, Yuhao Tan, Mingyao Pan, Hongyi Zhang, Bo Li

## Abstract

Mitochondrial (MT) mutations serve as natural genetic markers for inferring clonal relationships using single cell sequencing data. However, the fundamental challenge of MT mutation-based lineage tracing is automated identification of informative MT mutations. Here, we introduced an open-source computational algorithm called “MitoTracer”, which accurately identified clonally informative MT mutations and inferred evolutionary lineage from scRNA-seq or scATAC-seq samples. We benchmarked MitoTracer using the ground-truth experimental lineage sequencing data and demonstrated its superior performance over the existing methods measured by high sensitivity and specificity. MitoTracer is compatible with multiple single cell sequencing platforms. Its application to a cancer evolution dataset revealed the genes related to primary BRAF-inhibitor resistance from scRNA-seq data of BRAF-mutated cancer cells. Overall, our work provided a valuable tool for capturing real informative MT mutations and tracing the lineages among cells.

**Teaser:** MitoTracer enables automatically and accurately discover informative mitochondrial mutations for lineage tracing.

## Introduction

Recent advances in single cell RNA/DNA sequencing have led to deeper understanding of the heterogeneous human cell populations (1). Such information enables dissection of the tumor microenvironment and recovery of cell lineages (2). Previous studies have shown that single cell sequencing technologies can detect naturally occurring somatic mutations, which act as natural cell barcodes for different clones and lineages within organisms, including single nucleotide variations (SNVs) and copy number variations (CNVs) (3). However, detection of single cell nuclear SNVs or CNVs by whole-genome sequencing is challenging due to high error rates and potential transcript end biases.

Compared with nuclear genome, the 16.6-kb long mitochondrial (MT) genome is small for cost-effective sequencing (4). Furthermore, mitochondrial genomes have large number of copies and higher mutation rate, which is estimated to be 10- to 100-fold higher than nuclear genomes (5). Mitochondrial variations can be detected by single cell RNA sequencing (scRNA-seq) and single cell assay for transposase-accessible chromatin-sequencing (scATAC-seq) (6). ATAC-seq is an ideal technology to capture the MT genome due to its complete openness. Unfortunately, the current protocol of scATAC-seq discarded the cytoplasmic contents, resulting in poor coverage of mitochondrial genome. Several powerful technologies have been developed to overcover this problem and to obtain high coverage sequencing data of mitochondrial genome, including MAESTER (7) and mtscATAC-seq (8).

Although these technologies provided the platform for high quality mitochondrial genomic data generation, mutation-based lineage tracing remains challenging. Specifically, there lacks a computational method that can automatically and accurately identify informative mitochondrial mutations to discriminate cellular lineages. Several lineage reconstruction methods are available for single cell RNA/DNA sequencing, such as mgatk (9), SClineager (10), maegatk (7) and MQuad (11). The first three methods are developed specifically for scATAC-seq or scRNA-seq. The last method could be used across different single cell sequencing assays, but the accuracy of clonal inference in real human data remains unsatisfactory.

To lift these limitations, we develop a new software, MitoTracer, a complete and automatic computational method for informative mitochondrial mutation identification from scRNA or scATAC-seq samples. This pipeline performs all the necessary analysis steps, including mapping reads, generating mitochondrial variant allele frequency matrix, selecting informative mitochondrial mutations, and inferring potential clonal structures. We evaluated MitoTracer using three gold-standard datasets sequenced by bulk ATAC-seq, scRNA-seq and scATAC-seq. These results demonstrated that MitoTracer is a complete, highly sensitive, and efficient method for informative mitochondrial mutations that outperforms existing computational methods in scope and accuracy.

## Results

### MitoTracer is an automated pipeline for single cell lineage tracing using mitochondrial mutations

MitoTracer is a computational algorithm that identifies informative mitochondrial (MT) mutations for uncovering cell lineages or clones. It utilizes droplet-based (10X genomics) or plated-based (Smart-seq2) single cell sequencing data to detect MT mutations, such as 10X scRNA-seq data, 10X scATAC-seq data, modified 10X scRNA/DNA-seq data (MAESTER and mtscATAC-seq), and full-length Smart-seq2 data. We employed MERCI-mtSNP (*12*), a mutation-calling tool developed by our team, to identify MT RNA/DNA mutations and to obtain a variant allele frequency (VAF) matrix for each cell (Figure 1A). VAF may be impacted by various types of noise, such as sequencing errors and insufficient coverage. Compared to nuclear genome, the MT genome usually has significantly higher coverage due to shorter length. Therefore, controlling for sequencing errors is critical for detecting informative MT mutations. To solve this issue, we employed a previously described framework (*13*) to calculate the statistical power for mutation detection while adjusted for sequencing errors. We also eliminated mutations with extremely low or high cell-level frequencies, as such mutations are less useful to separate lineages or clones (Figure 1B, Step 1).

**Figure 1.**
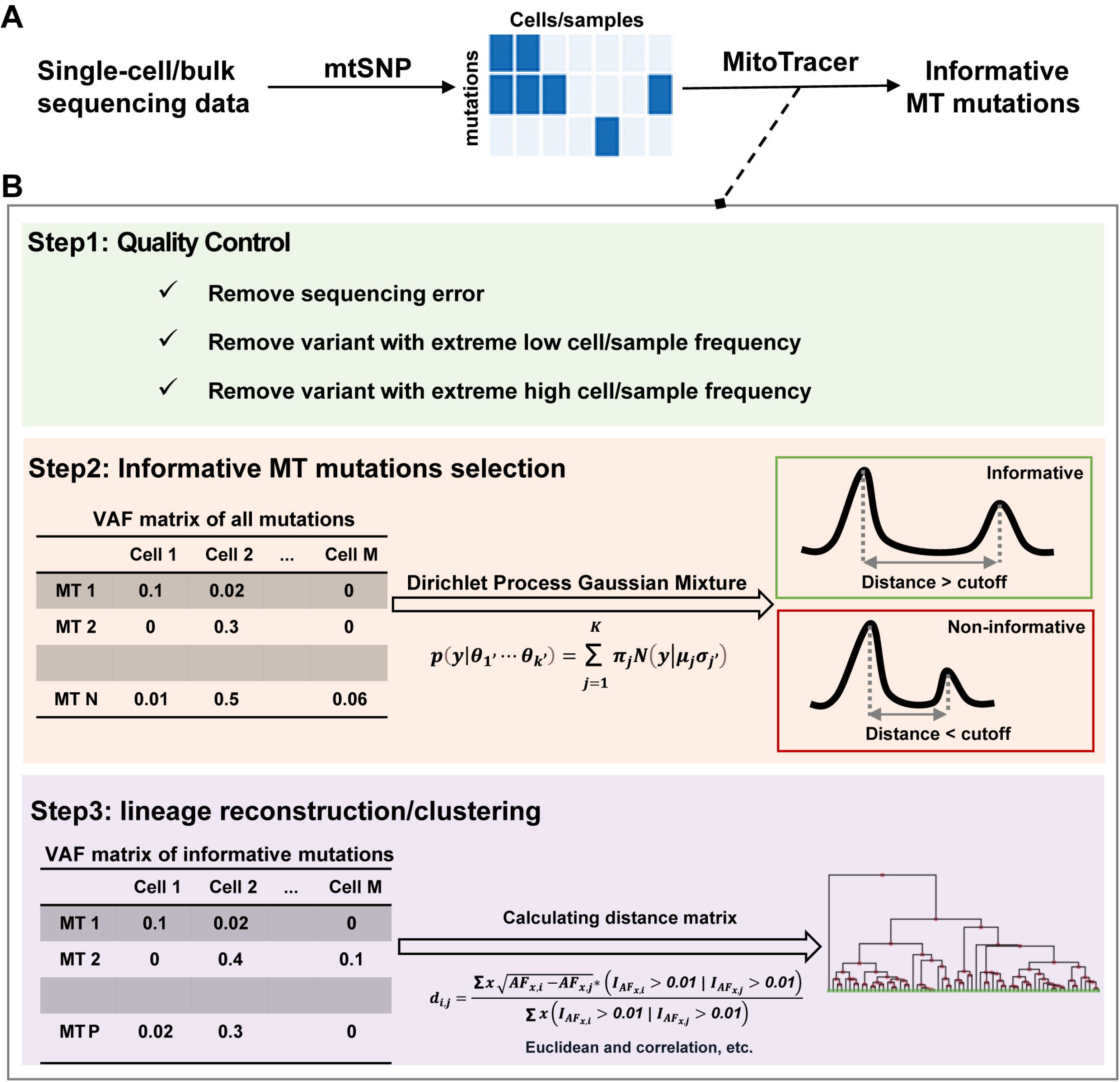
Overview of MitoTracer algorithm. (A) The whole analysis process of identifying informative MT mutations. MERCI-mtSNP calls MT mutations from single cell RNA or DNA sequencing data. The VAF matrix is generated for MitoTracer. (B) Informative MT mutation selection. MitoTracer firstly removes mutations caused by sequencing errors, and filters variants with extremely low/high cell- or sample-level frequency. Dirichlet process Gaussian mixture model is conducted on each MT mutation to find out informative MT mutation. We define the informative MT mutation as the absolute difference between the mean in the top two Gaussian distributions larger than the cutoff. MitoTracer uses the VAF matrix of these informative mutations to calculate the similarity matrix based on several distance methods, including the mitochondrial distance defined by ourselves, Euclidean, and correlation.

MitoTracer introduced a feature-selection process that automatically selects highly informative mutations for distinguishing lineages using single cell sequencing data. For each mutation, the VAF vector (VAF values across all cells) is assumed to follow a Gaussian mixture distribution. The number of Gaussian peaks reflect the count of distinguishable clones in the sample, which is usually unknown in real-world scenario. Therefore, we applied Dirichlet Process (DP) prior, a commonly used technique to model Gaussian mixture distributions with unknown number of modes (*14*). Variants with at least two peaks identified from DP were kept for downstream analysis. We compared the positions of the top two distributions assigned by DP to prioritize potential informative MT mutations for lineage detection. Specifically, if the two peaks are located far away, it indicates that the two clones they represented are sufficiently distinct in the allele frequencies of the given variant, and therefore, the variant is considered informative. The cutoff of the distance between the top two peaks is user-defined, with a default value 0.05 (Figure 1B, Step 2). Next, MitoTracer generated a mitochondrial distance matrix (Materials and Methods) using the selected mutations and performed hierarchical clustering to infer the lineage relationships between different cells, displayed as heatmap with dendrogram (Figure 1B, Step 3). Further details regarding quality control, informative MT mutation selection, and lineage reconstruction can be found in the Materials and Methods section.

### MitoTracer achieves better performance of lineage tracing

MitoTracer was benchmarked using data generated from cell lines with known lineage relationships. Specifically, human leukemia TF1 cells were sequentially cultured, with the initiation of next generation being a subclone of the current cells. Multiple subclones were cultured to acquire siblings of each generation. To capture the entire mitochondrial genome with high coverage, both the original and expanded clones were profiled using bulk ATAC-seq (Table S1) (9).

Using the above dataset as golden standard, we compared the accuracy of lineage reconstruction of MitoTracer and three other approaches, including informative MT mutations called at ≥ 0.2 (GTEx heteroplasmy threshold) (9), MQuad (scRNA-seq or scDNA-seq) (11), and SClineager (scRNA-seq) (10). The dataset comprised of 65 cell populations from 15 clones (A9 to G11). Overall, VAF_cutoff, MitoTracer, MQuad, and SClineager identified 57, 22, 1076, and 19 informative MT mutations, respectively (Table S2). In the original study, 44 MT mutations (Table S3) were manually selected as informative markers to reconstruct the experimental lineage (9). Of the four methods compared, the numbers of the selected mutations that overlapped with the manual set were 6, 15, 17, and 9 for VAF_cutoff, MitoTracer, MQuad, and SClineager, respectively (Table S4). Notably, MitoTracer had the highest percentage of overlap (15 out of 22), suggesting MitoTracer can specifically recognize lineage-associated MT variants with high sensitivity. We next evaluated lineage separation accuracy of each method by using the variants it selected. MitoTracer showed superior performance compared to the other strategies by correctly assigning all the cell populations into their corresponding clones (Figure 2A-2D). Hierarchical clustering performed on individual samples using MitoTracer grouped most of the subclones to the right parental clone (Figure 2B).

**Figure 2.**
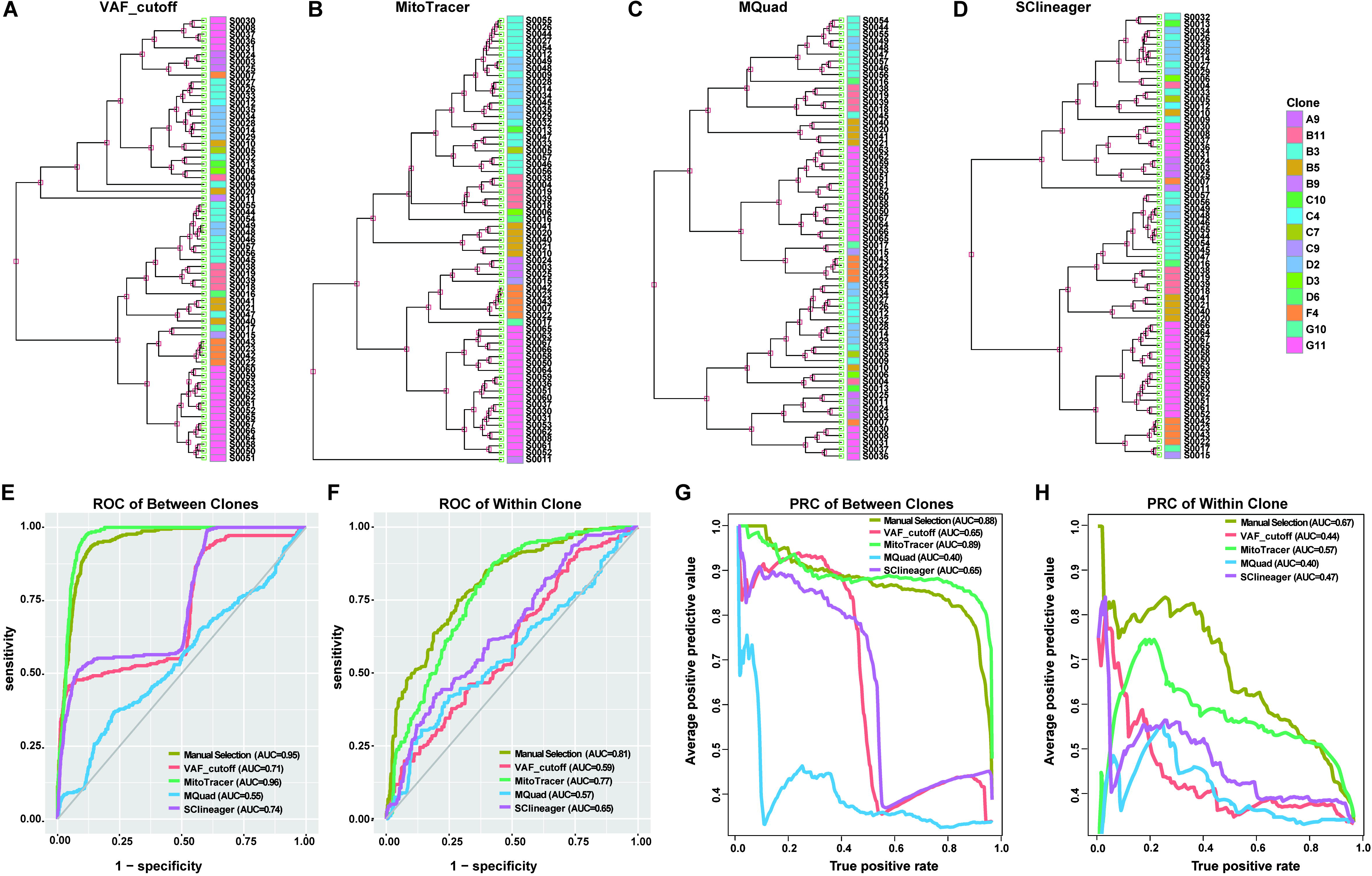
Performance comparison. (A-D) Performance comparison on a gold-standard dataset with 15 clones among four methods, including (A) VAF_cutoff, (B) MitoTracer, (C) MQuad, and (D) SClineager. Clone information is labeled by the “Clone” annotation bar on the right side of the heatmap. (E-F) ROC of the manual selection method and the above four methods under between-clone and within-clone. (G-H) PRC of the manual selection method and the above four methods under between-clone and within-clone.

To evaluate the accuracy of MT mutation-based fine-scale lineage reconstruction, we utilized a previously employed approach (9). Specifically, given any set of 3 samples where a true pair of siblings exists, each method is tested for the ability to select the siblings out of the sample triplet. Siblings are defined as the clones that share the same parental clone, thus the ‘Most Recent Common Ancestor’, or MRCA. Subsequently, we calculated the MT distance for each sample pair using the VAF matrix of informative MT mutations within a triplet. As siblings share MRCA, their MT distance is expected to be smaller than the other distances. Within each triplet, we defined MT distance between each pair of samples as the predictor, and sibling status as the response (sibling=1, non-sibling=0). This setting allowed us to evaluate the prediction accuracies of all methods by setting continuous cutoffs on the predictor. We then calculated the area under the curve (AUC) for both the receiver operating characteristic (ROC) curve and the precision-recall (PR) curve. We further tested the performances under two scenarios: the non-sibling sample is derived from the same MRCA (within-clone) and from a different MRCA (between-clones). Within-clone sample is expected to be genetically ‘closer’ to the true sibling pair, and thus more difficult to distinguish from. In both scenarios, MitoTracer showed the best overall accuracies in identifying the true sibling pairs compared to the other three automated methods (Figure 2E-H). We included the ROC and PR curves for the manually selected 44 MT mutations as a standard for optimal performance, where in both scenarios, only MitoTracer achieved similar AUCs.

### Robust identification of informative mitochondrial mutations from scRNA-seq and scDNA-seq data

Next, we evaluated the utilities of MitoTracer on single cell datasets. We conducted an initial analysis on a set of 5,842 cells derived from the BT142 and K562 cell lines, comprised of 1,251 BT142 cells and 1,101 K562 cells, respectively (Table S1). This dataset was produced using the MAESTER platform, which is compatible with the 10x Genomics 3’ protocols (7). The MAESTER technology enhanced the coverage of MT genome by enriching all MT transcripts. We conducted unsupervised hierarchical clustering on the VAFs of 24 informative MT mutations called by MitoTracer. Our analysis revealed two distinct clusters, which perfectly aligned with the two cell lines (Figure 3A). Similarly, we examined a dataset with the same setting but generated with another MT sequencing technology mtscATAC-seq (8), which was comprised of 437 cells from the TF1 and 519 cells from the GM11906 cell line. MitoTracer called 47 informative MT mutations, using which all cells were partitioned into two distinct clusters, matching the cell line annotations (Figure 3B). This result demonstrated the ability of MitoTracer to faithfully recover lineages from different genetic backgrounds.

**Figure 3.**
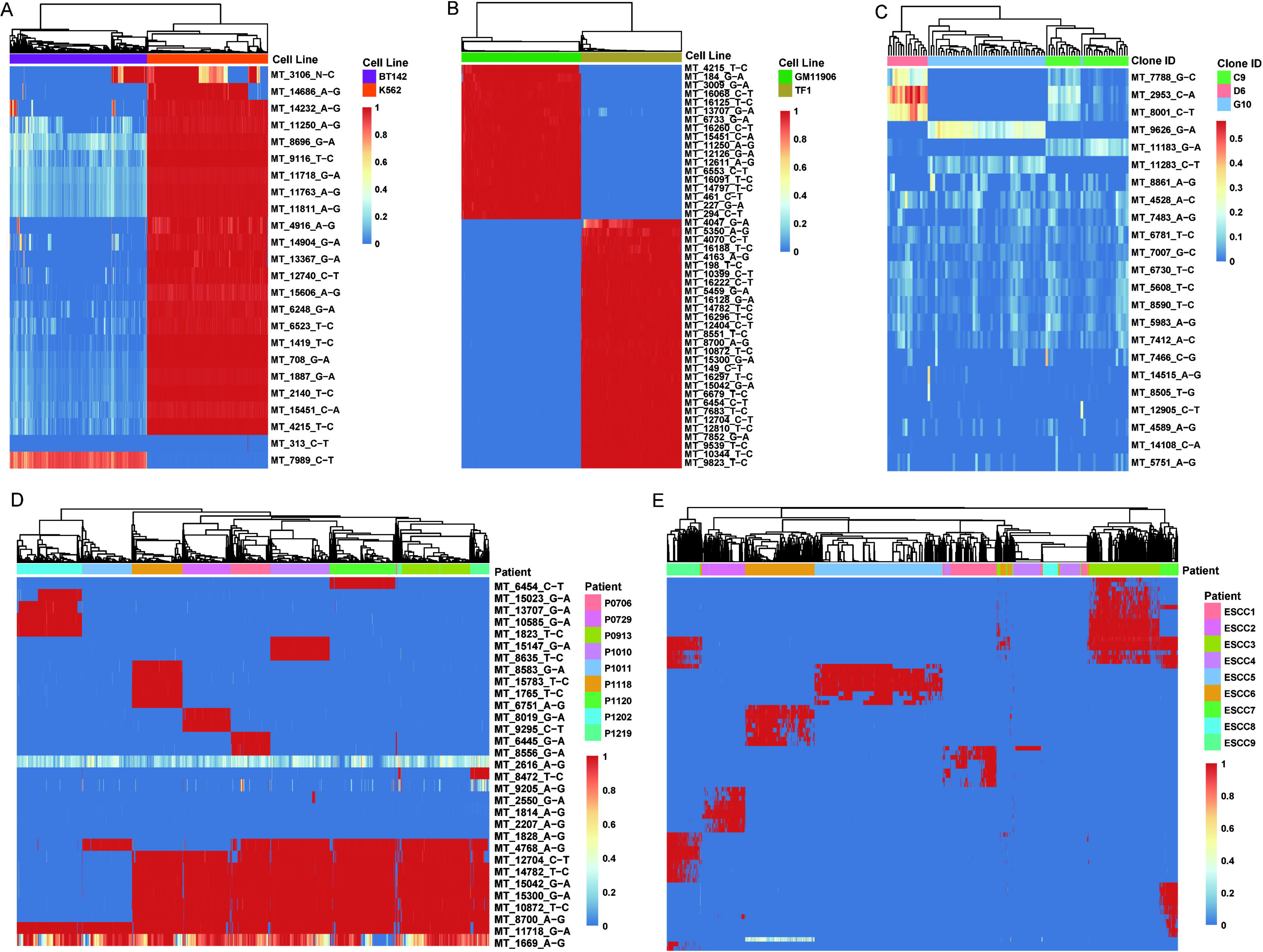
Validation of the MitoTracer algorithm. (A) Unsupervised hierarchical clustering of the VAF matrix of 24 informative MT mutations showed the clearing clustering of BT142 and K562 cells for the MAESTER dataset. (B) Unsupervised hierarchical clustering of the VAF matrix of 47 informative MT mutations showed the clearing clustering of GM11906 and TF1 cells for mtscATAC-seq dataset. (C) Unsupervised hierarchical clustering of the VAF matrix of 23 informative MT mutations showed the clearing clustering of C9, D6, and G10 hematopoietic cells for the scATAC-seq dataset. (D) Unsupervised hierarchical clustering of the VAF matrix of 31 informative MT mutations showed the clearing clustering of cells by their patient origin for the SMART-seq2 dataset. (E) Unsupervised hierarchical clustering of the VAF matrix of 82 informative MT mutations showed the clearing clustering of cells by their patient origin for the 10X scRNA-seq dataset.

We then tested the performance of MitoTracer using cells derived from the same genetic background. This dataset consisted of 96 TF1 cells profiled with scATAC-seq data, including three clones, C9 (n = 32), D6 (n = 16), and G10 (n = 48) (Table S1). These clones were defined based on culture history (9). MitoTracer identified 23 informative MT mutations to perform unsupervised clustering of all cells, which resulted in clearly separated clusters (Figure 3C). The three TF1 clones were accurately grouped into distinct branches of the TF1 dendrogram, demonstrating that MitoTracer can successfully recover the lineages of cells within the same genetic background.

The above analyses were performed using DNA samples. We next tested MitoTracer using RNA-seq samples, which is considered a more challenging task due to the relatively lower and uneven coverage in the mtDNA region. First, we applied MitoTracer to a Smart-seq2 dataset with 8,270 cells from 9 lung cancer patients (*15*) to evaluate its performance on cells with different genetic backgrounds (Table S1). Patient origins were utilized as class labels. MitoTracer called 31 informative MT mutations by pooling all the cells. Unsupervised clustering revealed that cells from different patients were grouped into separate groups, each with one or more representative MT mutations (Figure 3D). Despite the existence of 15 patient-specific MT mutations, MitoTracer successfully detected the common MT mutations shared by a subset of lung cancer patients. These findings demonstrated that MitoTracer is capable of accurately inferring genetic information from different cancer patients.

Last, we tested MitoTracer using the regular 10X scRNA-seq data, which is the most challenging task given the low coverage in the mtDNA region. We conducted the same analysis on a 10X scRNA-seq dataset of esophageal squamous cell carcinoma (ESCC), which contained 208,659 single cells from 60 individuals (*16*) (Table S1). We randomly selected 9 ESCC patients with a total 155,56 cells from this dataset, from which MitoTracer identified 82 informative MT mutations (Table S5). The number of patients matched that of the lung cancer dataset for comparison purpose. Hierarchical clustering revealed that almost 90% cells from different patients were separated into distinct groups (Figure 3E), thus confirming the ability of MitoTracer to detect patient-specific MT mutations from regular 10X scRNA-seq samples.

### Intrinsic BRAF inhibitor-resistant clones detected by MitoTracer

We next evaluated if MitoTracer could provide biological insights through the identification of lineage-specific MT mutations from cells of the same genetic background. Specifically, MitoTracer was tested on a Fluidigm scRNA-Seq dataset of 451Lu melanoma cells harboring the BRAF V600E mutation (*17*). This dataset contained 162 unselected parental cells and 157 cells resistant to BRAF inhibitors (Table S1). MitoTracer detected 24 informative mutations in mtDNA. Unsupervised clustering on the VAFs of these mutations uncovered two major clusters: Cluster 1 predominantly contained parental cells; Cluster 2 were mostly BRAF inhibitor-resistant cells (Figure 4A). Interestingly, we observed a sub-branch within Cluster 2 that contained 37 parental cells and 2 BRAF inhibitor-resistant cells. We postulated that this sub-branch represented an unselected clone with intrinsic resistance to BRAF inhibitor. Notably, the “MT_16389_G-A” mutation was exclusively detected in the resistant group (Cluster_2). The VAF of this mutation in the intrinsic clone was lower than that in the other BRAF inhibitor-resistant cells in Cluster 2, suggesting that this variant might be under positive selection (Figure 4B). Furthermore, we identified differentially expressed genes (DEGs) by comparing parental cells with and without the “MT_16389_G-A” mutation. We detected 647 genes that displayed significant changes (Table S6; Method was described in the Materials and Methods section). Notably, within the top 10 upregulated genes, seven are identified as having connections to drug resistance, including COX1 (*18*), COX2 (*19*), MT2A (*20*), FTH1 (*21*), MIF (*22*), MALAT1 (*23*), and CYTB (*24*). Furthermore, recent investigations have underscored the frequent upregulation of COX2 in a range of human cancers, including melanoma, colorectal, breast, stomach, lung, and pancreatic tumors (*25, 26*) Previous research has demonstrated the efficacy of COX2 inhibition in overcoming therapeutic resistance in BRAFV600E colorectal cancer (*27*) and its pivotal role in addressing drug resistance in melanoma (*19, 28, 29*).

**Figure 4.**
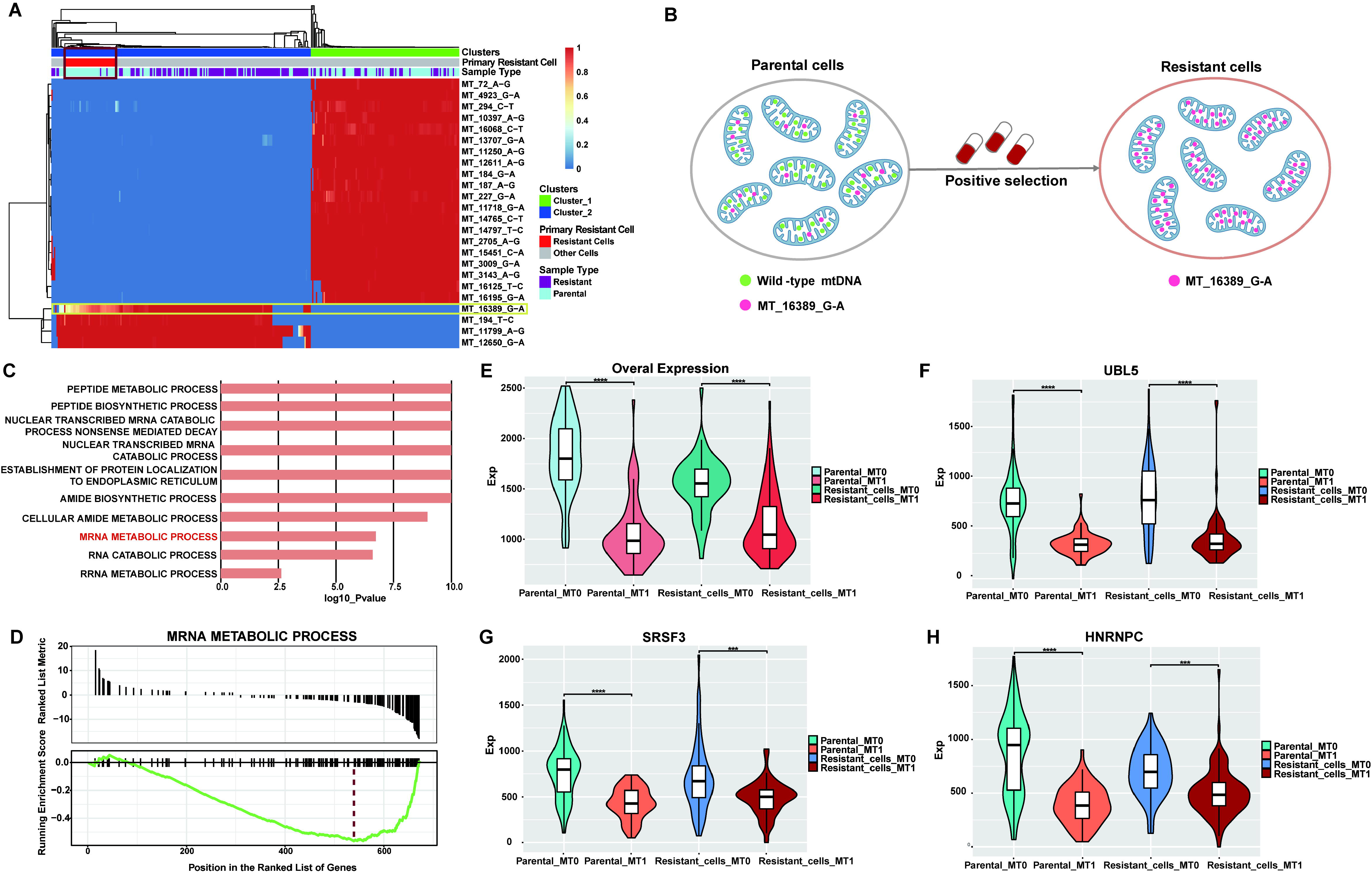
MitoTracer identified essential primary BRAF inhibitor-resistant genes. (A)The reconstructed lineage was visualized by heatmap from MitoTracer. All these cells were clustered into two major clusters labeled “Cluster_1” and “Cluster_2”. Cells were labeled according to their original BRAF inhibitor resistance status, “Resistant” and “Parental”. We also labeled the primary resistant cells predicted by MitoTracer with “Resistant Cells”. (B) Graphic description of positive selection for primary resistant-related MT mutation, MT_16389_G-A. (C) Gene ontology biological process enrichment results of 674 differentially expressed genes. (D) GSEA enrichment results of 674 differentially expressed genes. (E) The overall expression level across resistant and MT_16389_G-A status. We defined the group “MT0” which presented the cells with MT_16389_G-A mutation, and MT1 indicated the cells without MT_16389_G-A mutation. (F-G) the expression level of PSMB3, SNRPD2, and UBL5 across resistant and MT_16389_G-A status. The definition of the group was the same as (E). ***P<0.0001 and **** P<0.00001.

Gene Set Enrichment Analysis (GSEA) revealed that the top 647 DEGs were enriched in metabolic processes (Figure 4C), a relationship that has been established for BRAF inhibitor resistance (*30*). Notably, the down-regulated genes were significantly enriched in mRNA metabolic process (Figure 4D). We subsequently evaluated the overall expression levels of this gene set in parental/resistant cells with or without the “MT_16389_G-A” mutation. Interestingly, we found that both parental and resistant cells with the “MT_16389_G-A” mutation displayed significantly lower expression levels of genes related to this process than the other cells (Figure 4E). A few of such genes included: ubiquitin-like protein 5 (UBL5) (Figure 4F), Ser/Arg-rich splicing factor 3 (SRSF3) (Figure 4G), and heterogeneous nuclear ribonucleoprotein C1/C2 (HNRNPC) (Figure 4H). Previous studies have linked UBL5 to melanoma growth as deubiquitinating enzymes (DUBs) (*31*), while depletion of SRSF3 leads to a switch in MDM4 splicing that influences p53-mediated antiproliferative activity (*32, 33*). Dysregulation of HNRNPC has been observed in lung cancer, breast cancer, and oral squamous cell carcinoma patients (*34–36*). This gene has been shown to regulate tumor cell proliferation and promote radiation resistance in pancreatic cancer (*37*) and mediate mRNA stabilization to alter energy metabolism, facilitating metastasis and invasion of oral cancer cells (*38*). Collectively, these findings suggested that the top genes revealed by MitoTracer analysis may play a role in primary BRAF-inhibitor resistance and represent potential targets for reversing resistance.

## Discussion

Here, we developed a fully automated lineage tracing method by using the single cell sequencing data, eliminating the need of manual marker selection. Our benchmark analysis showed that MitoTracer could reliably identify informative MT mutations and reconstruct lineage from diverse single cell genomic data, including scATAC-seq, Smart-seq2, 10X scRNA-seq, and variants of 10X platform single cell data. We systematically validated our approach for both intra- and inter-patient lineage reconstruction and demonstrate its capability of deriving biological insights.

Lineage tracing based on mitochondrial mutations has provided important insights through recent analysis of patient samples. Zhang et al. applied this technique to construct phylogenetic trees for MKI67+ T cells and macrophages by using the scRNA-seq data derived from hepatocellular carcinoma patient samples (*39*). Their findings illuminated shared lineages between cells in the tumor and ascites, suggesting the plausible origin of subsets of lymphocytes and macrophages in ascites from the tumor. In another study, Wang et al. delineated the mesenchymal-to-proneural hierarchy from glioma stem-like cells (GSC) through mitochondrial mutations in glioma. This observation substantiates the role of mesenchymal GSCs as the progenitors of proneural GSCs (*40*). Collectively, these applications underscored the effectiveness of lineage tracing via mitochondrial mutations as a powerful technique to trace cell migration patterns and to reveal the lineage relationships among stem-like cells in malignant tumors.

A similar observation in this work as seen in the previous studies is that despite the high mutation rate of mitochondrial genome, informative variants within a subject remain few. Previous study has suggested an ~10-fold higher rate in mitochondrial DNA than in nuclear DNA (*41*). The mutation rate of nuclear genome is estimated to be 0.06 × 10^−9^ per site per cell division (*42*), and thus for mitochondrial genome, this rate is 0.06 × 10^−8^. We assume each cell contain at least 100 copies of mtDNA, and estimated the per division mutation rate of mitochondrial genome to be approximately 0.001. Although this is a very high rate, after 30 cell divisions, the expected number of cells carrying a mtDNA mutation is approximately 2.3 × 10^7^ (*43*), which account for only 2% of the population. Statistically speaking, a variant ideally should have high heteroplasmy (within cell variant frequency) and high intercellular variation. These criteria required the variant to occur early enough for population fixation and segregation, which usually need more cell divisions. Hence, tracing of finer lineages, such as lymphocyte clonal expansion and differentiation upon antigen recognition, remains a computational challenge.

There are also limitations of MitoTracer. First, the application of our approach to 10X data presents challenges arising from uneven and low coverage. Another limitation is that the sensitivity to detect smaller clones is anticipated to be suboptimal. Therefore, an accurate and deep sequencing of mitochondrial genome is required for detecting clone-specific MT mutations by MitoTracer. Finally, most of our conclusions are of an exploratory nature and lack validation through additional experimental evidence. Although the biological effects of most mitochondrial mutations investigated in this context remain uncertain, the precise identification of these mutations and the elucidation of their biological functions are crucial avenues for further exploration.

Overall, the amount of scRNA-seq data in the public domain had significantly increased in recent years, however, the algorithms for mining these datasets were still limited, especially for the MT genome. Lineage tracing by informative MT mutations is a powerful approach to reach this goal. Thus, MitoTracer is likely to be broadly useful and immediately applicable, because it can automatically and accurately identify the informative mitochondrial mutations for lineage tracing and better understanding the biological processes from an alternative angle.

## Supporting information

All public datasets used in this study.

The list of MT variants called by four methods.

The 44 MT variants selected by Vijay.

The MT variants overlap between manual set and four methods.

The list MT variants called by MitoTracer in ESCC patients.

The 647 DEGs list.

## Acknowledgments

This work is supported by NCI 1R01CA245318 (B.L.), 1R01CA258524 (B.L.), and NCI P50 CA070907 (JDM, HL).

## Data availability

All datasets used in this study are publicly available in GEO, EGA or SRA. The detailed information for these datasets was shown in Table S1.

## Code availability

The source code of MitoTracer R package, and the codes to run example data are available at: https://github.com/xuexinyu11/MitoTracer. Any additional information is also available from the corresponding author upon request.

## Methods

### MT mutation detection algorithm MERCI-mtSNP

Zhang et al. developed MERCI-mtSNP (*12*) for calling single-nucleotide variants (SNV) in MT genomics data generated from popular bulk or single cell sequencing technologies, such as 10x Genomics scRNA-seq, scATAC-seq, smart-seq2 and bulk ATAC-seq. The aligned bam file was used as the input of MERCI-mtSNP and extracted all reads aligned to the MT genome. For 10X single cell sequencing data, all reads were separated by cell barcodes and generated new MT bam file only containing the extracted MT reads. MERCI-mtSNP called MT variants for each cell with at least K mitochondrial reads (K = 1,000 for scRNA-seq data, K = 2,000 for scATAC-seq data). And then, the VAF for each altered base at a given locus is the number of the supporting reads divided by the total read depth. In order to get high-quality variants, MERCI-mtSNP used the reads with base-quality score (base-quality score >15 for scRNA-seq data, base-quality score >25 for scATAC-seq data) to calculate the VAF values. At last, one csv file and one txt file would be generated to represent the information of MT coverage and variants, respectively.

### MT mutations sequencing error filtering

To remove mutations that resulted in sequencing errors, we implemented additional filtering because its comparatively higher coverage per site per cell can result in more sequencing errors. Given the random sequencing error rate *e*, the probability of observing at least m identical alternate reads due to sequencing error can be represented as:

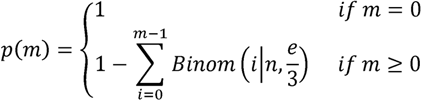

We then calculate the minimum number of alternate reads k supporting that the P(k) is less than a defined false-positive rate (FPR):

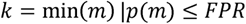

We specified the sequencing error rate *e* = 1 × 10^−3^ and *FPR*=5 × 10^−7^ as the default values in this study.

### MitoTracer model

We developed a Dirichlet process Gaussian mixture model to identify the clone or lineage-specific MT mutations. The informative MT mutations are heteroplasmic and only mutate in specific sub-cell populations. We hypothesized that the VAF distribution of the real informative MT mutations was a mixture of several Gaussian distributions. Thus, we can use the Dirichlet process to dissect the number of Gaussian distributions and estimate the densities for each MT mutation. If *y_i_*denotes the ith row in VAF matrix with N MT mutations (rows) and M cells (columns), we rescale *y_i_* such that its mean is 0 and the standard deviation is 1, then the model is:

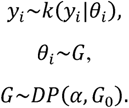

Where 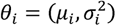 denotes the mean and variance, and *G_0_* is the base measure.

Rescaling y_i_ leads to the default parameterization of *G_0_* being uninformative. If we assume the Gaussian mixture model has *K* components, this model may be written as:

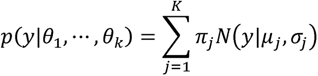

Where *θ_j_* = {*μ_j_*, *σ_j_*, *π_j_*) is the set of parameters for component *j*, *π* are the mixing proportions or weights (which must be positive and sum to one). We used Markov Chain Monte Carlo (MCMC) algorithms for inference on this model. The Markov chain relies on Gibbs updates, where each parameter is updated in turn by sampling from its posterior distribution conditional on all other parameters. We repeat this process for 10,000 times when the cell number is smaller than 100. In general, the total iteration of 2,000-5,000 should be sufficient.

We ordered mixture Gaussian distributions by mixing proportions for each MT mutation and calculated the mean difference between top two distributions. The MT mutation with the larger difference indicates more informative.

Finally, the mean difference cutoff for selection of informative mutations could be set manually, with a default value of 0.05.

### Distance matrix of cells or clones

The distance matrix D is the matrix whose entries are the pairwise distances between clones or cells. We define D for pairs of observations *i, j* over informative MT mutations (*x*) by the allele frequency matrix, only MT mutations with sufficient allele frequency in at least one clone or cell are included (minimum allele frequency > 0.01). We define the distance matrix between observations *i,j* by the distance *d_i,j_* as follows:

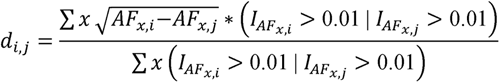

Where *I* is the indicator function.

### Data source, processing and read alignment

The comprehensive information for all datasets utilized in this paper has been provided in Table S1. Raw fastq files for public data were downloaded from Gene expression Omnibus (GEO), European Genome-Phenome Archive (EGA) and Sequence Read Archive (SRA).

For each library, raw fastq files were aligned using either Bowtie2 (bulk ATAC-seq) (*44*), STAR version 2.7.2b (SMART-seq and Fluidigm scRNA-Seq) (*45*), Cell Ranger (V6.0.0) Software Suite (10X single cell RNA/DNA sequencing data) to the GRCh38 reference genome. All the output bam files were utilized for mitochondrial variants calling by MERCI-mtSNP.

### Differential gene expression analyses

After normalization, the data matrix contained 34,806 genes. Significantly differentially expressed genes were identified using the eBayes function in limma R package, comparing parental cells contained “MT_16389_G-A” mutation with wild-type parental cells as the baseline. We adjusted p values (q values) for multiple testing using the Benjamini–Hochberg method. The differentially expressed genes (q < 0.01) with log2 fold change (FC) > 1 were identified as upregulated genes, while those with log2 fold change (FC) < 1 were identified as downregulated genes. All differentially expressed genes were ranked by log2 fold change. Pathway enrichment was performed on ranked lists with gene set enrichment analysis (GSEA) using KEGG and Gene Ontology.

### Statistical Analysis

Computational and statistical analyses in this work were performed using the R programming language v4.2.3. FDR control was using the Benjamini-Hochberg method. ROC curves, PR curves, and AUC values were generated using package ROCR (v1.0-11). Heatmaps were generated using R package pheatmap (v1.0.12). Differentially expressed genes were identified by R package limma (v3.54.2). GSEA was performed by R package msigdbr (v7.5.1) and clusterProfiler (v 4.10.0). Subpanels of main figures were produced using ggplot2 (v3.4.2).

## Supplementary Tables

**Table S1. All public datasets used in this study.**

**Table S2. The list of MT variants called by four methods.**

**Table S3. The 44 MT variants selected by Vijay.**

**Table S4. The MT variants overlap between manual set and four methods.**

**Table S5. The list MT variants called by MitoTracer in ESCC patients.**

**Table S6. The 647 DEGs list.**

